# Genome assembly of *Bougainvillia* cf. *muscus* (Cnidaria: Hydrozoa)

**DOI:** 10.1101/2025.01.24.634790

**Authors:** Aide Macias-Muñoz, Rebecca Varney, Eva Katcher, Maia Everhart, Todd H. Oakley

## Abstract

**Background:** As one of just a handful of non-Bilaterian animal phyla, Cnidaria are key to understanding genome evolution across Metazoa. Despite their importance and diversity, the genomes of most species in the phylum are unsequenced, due in large part to difficulties cultivating them in a laboratory. Here, we present a genome sequence of *Bougainvillia* cf. *mucus*, a hydrozoan with four marginal bulbs each containing seven simple eyes (ocelli). This species appeared in our tanks from contamination. While we lacked sufficient samples for transcriptomic or functional studies, we were able to expand our knowledge of how the genome of this species compares to the few, better studied members of hydrozoans by investigating synteny to other cnidarians, repetitive element content, and phylogenetics and synteny of vision-related genes in this eyed species compared to eyeless relatives.

**Results:** The genome sequence consists of 350 contigs with an N50 of 10 Mb, a total genome length of 375.328 Mb, a BUSCO score of 90.1%, and predicted protein coding genes totaling 46,431. We found a high degree of macrosynteny conservation with *Hydra vulgaris* and *Hydractinia symbiolongicarpus*. Repetitive elements make up 62% of this *Bougainvillia* genome. For vision-related genes, we identified 20 cnidarian opsins (cnidops) in *Bougainvillia* and found instances of gene duplication and loss in families associated with bilaterian eye development, phototransduction, and visual cycling.

**Conclusions:** This high-quality, contiguous genome in an eyed Hydrozoan will be a valuable resource for additional comparative genomic studies.

## Introduction

Comparative genomic studies are crucial for understanding the genetic underpinnings of biodiversity, phylogenetic relationships, genome evolution, and speciation. With high-quality genome assemblies, we can investigate gene duplications and losses, differences in genome size, conservation and divergence of synteny, and comparisons of *cis*-regulatory regions across a variety of clades. With broader sampling we can begin to address unanswered questions about the contribution of genomic changes to animal evolution. For example, chromosome-level assemblies for ctenophores and sponges coupled with a synteny analysis supported ctenophores as the sister group to other animals, clarifying a long debate (Schultz et al. 2023). Synteny analyses have also identified genomic changes correlated to major events in vertebrate evolution and a transition from marine to freshwater evolution of annelids (Simakov et al. 2020; Lewin et al. 2024). These recent studies highlight how new genomes can provide insight into the genetic changes correlated with animal evolution and diversity.

Comparative genomic studies have been impactful to our understanding of animal evolution. In Cnidaria, the phylum that encompasses jellyfish, sea anemones, and corals, genetics have been used to investigate cell differentiation, axis patterning, development, and regeneration (Broun & Bode 2002; Sebé-Pedrós, Chomsky, et al. 2018; Cazet et al. 2023) (see reviews Galliot & Schmid 2002; Technau & Steele 2012) These studies sought to determine whether gene function and gene networks were conserved between cnidarians and bilaterians. Since Cnidaria is one of the sister groups to Bilateria, comparative studies provide insight into genes that were present before their divergence. Cnidaria consists of ∼13,300 described species with unique morphologies, life histories, but so far only a few are represented in genomic studies (Kayal et al. 2018). For model cnidarians like *Nematostella, Hydra*, and *Hydractinia*, publicly available chromosome-level assemblies, transcriptomics, single-cell sequencing data, and transgenics have facilitated their use in a broad range of interdisciplinary research (Chapman et al. 2010; Sanders et al. 2018; Sebé-Pedrós et al. 2018; Siebert et al. 2019; Cazet et al. 2023; Zimmermann et al. 2023; Schnitzler et al. 2024; Salamanca-Díaz et al. 2024). Recent studies expanded the resources available for select cnidarians to include several high-quality genomes from jellyfishes (hydrozoans, cubozoans, and scyphozoans) and corals (anthozoans) (see review: Santander et al. 2022). However, model organisms make up the vast majority of genomic resources for cnidaria (12/44 available genomes in Hydrozoa are *Hydra*), therefore neglecting much of the vast diversity in form and function of the group. This is partially due to challenges in sampling and identifying clear, often small, gelatinous animals with few defining features (Daly et al. 2007). Improving this number might help identify some of the gene-level differences across cnidarian species that contribute to their diversity in form and function (Travert et al. 2023).

Cnidarians provide important study organisms for visual system evolution. Within the phylum, eyes evolved convergently at least 9 times (Picciani et al. 2018; Miranda & Collins 2019). Some Cnidarian eyes share expression of some gene families in common with the eyes of Bilaterian animals, although often expressing different specific orthologs (Plachetzki et al. 2007; Suga et al. 2008; Koyanagi et al. 2008; Vöcking et al. 2022). The eyes of one species in particular, the box jellyfish *Tripedalia cystophora,* are well studied, with published work on morphology, gene expression, light sensitivity, and visual behavior (Nilsson et al. 2005; Coates et al. 2006; Garm et al. 2007, 2011; Liegertová et al. 2015; Bielecki et al. 2014; Garm et al. 2022). To build on the diverse studies of vision in *T. cystophora*, we sought to sequence and assemble its genome. However, our culture of *T. cystophora* polyps unexpectedly began producing small hydrozoan medusae. At first, we assumed these small medusae were juvenile *T. cystophora* and sent them for genome sequencing before discovering they were a contaminant of the hydrozoan genus *Bougainvillia cf. muscus* (from hereon referred to as *Bougainvillia*). Unfortunately, we were never able to obtain polyps or additional medusae of the *Bougainvillia* contaminant, thus preventing further analyses such as gene expression. Although encountering one of the challenges in working with small cnidarians, we still were able to assemble a genome for an eyed jellyfish species that can be used as a representative for the family Bougainvilliidae within Filifera.

*Bougainvillia* belongs to a clade called Filifera, which besides some notable exceptions, have very few genomic resources (Kayal et al. 2015) The Filifera are a large suborder of the cnidarian class Hydrozoa that include the model *Hydractinia*, which lack eyes, along with the “immortal jellyfish” *Turritopsis*, which have eyes. Within Filifera, eyes probably evolved at least twice and potentially up to five times convergently (Picciani et al. 2018). This estimate is a range because of phylogenetic uncertainty., Picciani et al. accounted for this uncertainty in state transitions by naming uncertain nodes with letters and using numbers only for more clear transitions to eyed species (see Picciani et al. 2018, Fig.S1). Filifera had two convergent eye origins named eye origin 4 and 5, and 4a-d were additional possible convergent eye origins within the suborder. *Bougainvillia mucus* is representative of eye origin 4a, other *Bougainvillia* fall under eye origin 4b, *Nemopsis bachei* is representative of eye origin 4c, and *Turritopsis* represent eye origin 4d (Picciani et al. 2018). Based on this analysis, *Bougainvillia mucus* are representative of an eye origin independent from other eyed species in the genus. Future studies into the morphology and phylogenetic placement of species in this group could resolve the number and timing of the eye origins in Filifera.

Here, we present the sequenced genome of an eyed cnidarian from genus *Bougainvillia* (Fig 1). We generated a high-quality genome of 350 contigs across a total length of 375 Mb. This is the best genetic resource to date for this genus and family; most research thus far focused on morphological descriptions and distributions (Denitto et al. 2007; Nogueira et al. 2013; Batistić & Garić 2016; Mendoza-Becerril et al. 2018). We used this genome to place *Bougainvillia* in a species-level phylogeny of Cnidaria, to compare synteny in Hydrozoans, and to examine the placement and copy number of genes known to play roles in eye development and vision. This reference genome contributes additional taxon sampling to an understudied clade of Cnidarian, and we are confident it will provide a resource for future comparative studies.

**Figure 1.**
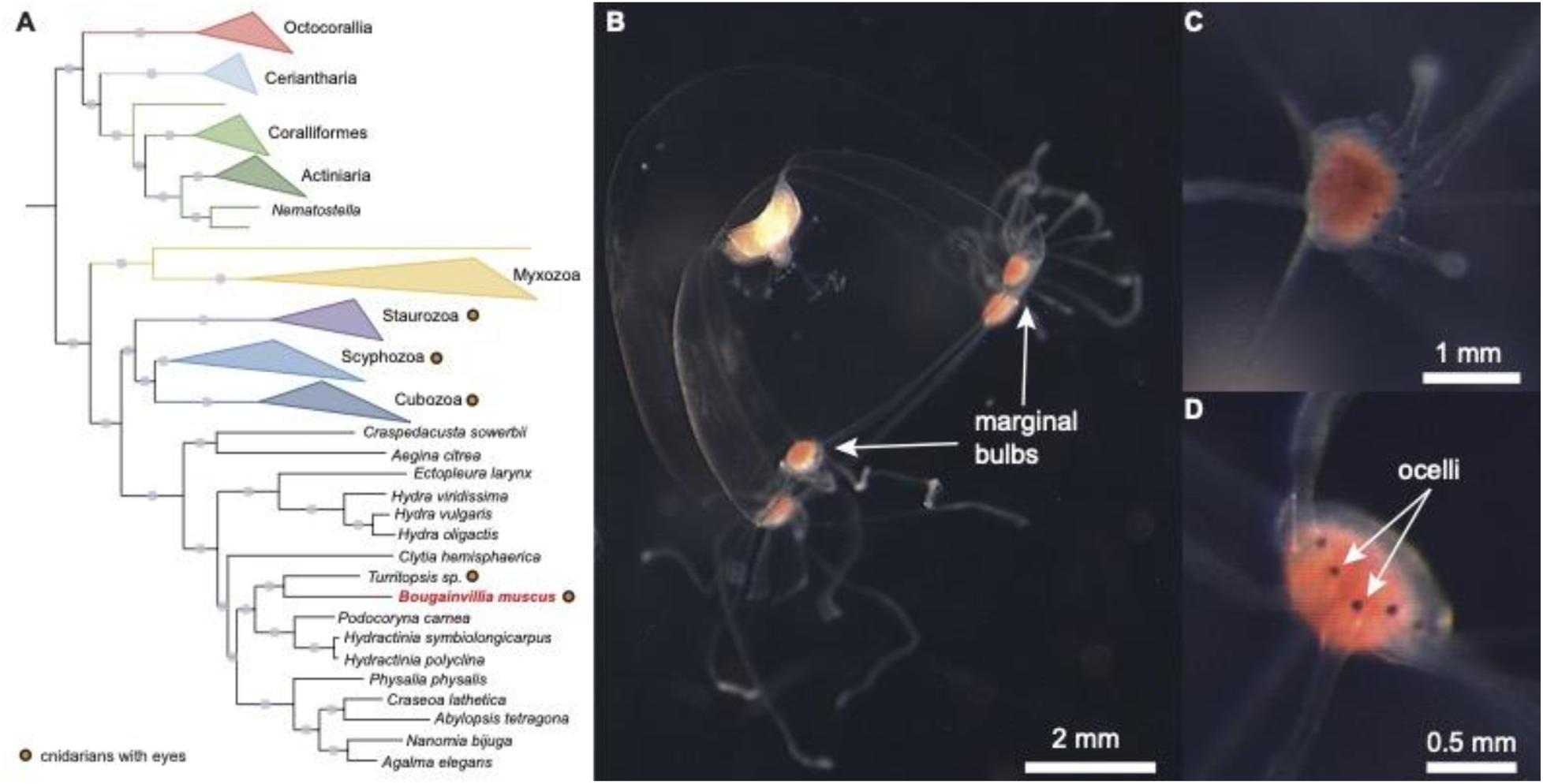
*Bougainvillia* cf. *muscus* falls within Hydrozoa and has ocelli. A) Cnidarian phylogeny using 748 orthologs from DeBiasse et al. 2023 with *Bougainvillia* cf. *muscus* added. Branches that have over 80% bootstrap support are labeled with light gray circles. Orange circles adjacent to species or class names represent cnidarians that have eyes. B) Photograph of a *Bougainvillia* medusa with marginal bulbs labeled. C-D) Zoomed in images of the marginal bulbs showing the ocelli which are rounded, black, and located at the base of each tentacle. *Bougainvillia* have 7 ocelli in each of their four marginal bulbs.

## Methods

### DNA extraction

We obtained *Bougainvillia* from a tank of seawater at 25℃ maintained at the University of California, Santa Barbara that also contained *Tripedalia cystophora* polyps. Water came from UCSB’s seawater system and the *T. cystophora* polyps came from a culture from Denmark, with *T. cystophora* originating from Puerto Rico or Florida. In September 2021, we began noticing many small medusae. We collected fifteen free swimming medusae using plastic pipettes and washed them thoroughly with filtered seawater. We extracted DNA from fresh tissue using Circulomics Nanobind Tissue Big DNA Kit (NB-900-701-01; Circulomics via Pacific Biosciences, Menlo Park, CA, USA) skipping the tissueRuptor step.

### Data processing

The UCI Genomics High Throughput Facility made PacBio HiFi low input libraries from our extracted DNA and sequenced them on 2 SMRT cells. We processed HiFi reads from BAM files using ccs v 6.4.0. In cell 1, 2,085,669 (46.63%) zero-mode waveguides (ZMWs) passed filter and 1,982,604 (48.63%) in cell 2. We used Bam2fastq v. 1.3.0 to combine the sequencing data from the two cells and to extract it as fastq files. We generated a genome assembly using hifiasm (Cheng et al. 2021), then determined completeness and contiguity using BUSCO v. 5.3.2 -l metazoa and BBmap v. 38.96 (Bushnell B. et al. 2014; Simão et al. 2015). To check for potential contaminants, we used blobtoolkit v. 4.2.1 and removed outlier contigs with no overlap to Cnidaria and with GC content <0.3 and >0.5 (Challis et al. 2020). We used BRAKER3 (Gabriel et al. 2024) to annotate the final assembly using protein models from *Hydra vulgaris* (Chapman et al. 2010), *Clytia hemisphaerica* (Leclère et al. 2019), *Hydractinia echinata* and *Hydractinia symbiolongicarpus* (Schnitzler et al. 2024)as input.

### Species identification

To maintain a record of the collected samples, we photographed medusae concurrently with DNA extraction. From the photographs, a colleague later suggested the medusae were *Bougainvillia*. We based an initial species identification on morphological characteristics by referencing World Register of Marine Species (WoRMS Editorial Board 2024)After assembling the genome, we extracted sequences for COI, 18S, 16S and 28S and compared them to available sequences in NCBI via BLAST. The top hit was *Bougainvillia muscus*. We then downloaded sequences of *B. muscus* COI, 18S, 16S and 28S from NCBI and did reciprocal BLAST+ to the assembled genome. The sequences had an e-value of 0 suggesting a very close, if not exact match. For further validation, we downloaded eleven 16S sequences for eight available *Bougainvillia* species from NCBI and generated a 16S phylogeny.

### Cnidarian species phylogeny

We obtained sequences for 748 genes previously aligned across cnidarians from (DeBiasse et al. 2024). We used hmm2aln.pl (https://github.com/josephryan/hmm2aln.pl) to identify sequences that matched each of these alignments in *Bougainvillia*. We then filtered for the best match (longest sequence with least gaps), ran MAFFT v. 7.520 (default parameters) and Gblocks v0.91b (-b2=10 -b3=10 -b4=5 -b5=a), and concatenated the matrix (see data availability). We generated a phylogenetic tree from the concatenated matrix using iqtree2 v. 2.2.2.6 -nt AUTO -bb 1000 -m TEST, and then used iTOL v6 to visualize, root (outgroups bilateria, ctenophora, and porifera), and annotate the tree. We also generated a tree from concatenated alignments prior to Gblocks using the same model.

### Macrosynteny and repetitive sequences

To characterize synteny conservation across Hydrozoans, we investigated macrosynteny between our final *Bougainvillia* genome assembly and the *Hydra* and *Hydractinia* chromosome-level genome assemblies (Simakov et al. 2020; Schultz et al. 2023; Kon-Nanjo et al. 2023). We downloaded the *Hydra vulgaris* v3 (GCF_022113875.1) and *Hydractinia symbiolongicarpus* v2.0 (https://doi.org/10.6084/m9.figshare.22126232.v1) genomes from NCBI and Figshare, respectively. We used odp with a minimum scaffold size of 1 Mb for both comparisons (Schultz et al. 2023). To estimate the degree of macrosynteny conservation, we calculated the number of one-to-one orthologs in homologous locations and divided by the total number of orthologs in the odp map (Wang et al. 2017; Li et al. 2022).

To identify repetitive sequences, we first generated a custom library using default parameters in RepeatModeler v. 2.0.5 (Flynn et al. 2020), then used it to identify repeats in the genome assembly with RepeatMasker v. 4.1.5 (http://www.repeatmasker.org).

### Vision-related phylogenetic trees

To investigate the phylogenetic relationships of vision-related genes across cnidarians, we downloaded publicly available transcriptomes from NCBI using NCBI’s Datasets command-line tools (Supplementary Table S1). We used transdecoder v. 5.7.1 to extract coding and peptide sequences and cd-hit v. 4.8.1 with parameters -c 0.9 -n 5 to remove redundant transcripts (Li & Godzik 2006; Fu et al. 2012). We used orthofinder v. 2.5.5 to identify orthogroups among all datasets (Emms & Kelly 2019). We aligned sequences for candidate genes using MAFFT v. 7.520 and generated trees using iqtree2 v. 2.2.2.6 -m MFP -B 1000 -alrt 1000 -T 8 (Katoh & Standley 2013; Minh et al. 2020). We used iTOL v6 to annotate our final tree files, rooted at midpoint (Letunic & Bork 2024). For the opsin tree, we downloaded opsin sequences from (McCulloch et al. 2023) and searched for similar sequences in the *Bougainvillia* genome assembly using command-line BLAST+. We extracted sequences for the top hits and searched against the NCBI database to confirm opsin annotation. We added all *Bougainvillia* sequences that matched an opsin annotation to the opsin fasta file from (McCulloch et al. 2023) and used them as input for MAFFT and iqtree2. We rooted the tree using *Trichoplax* placopsins (Feuda et al. 2012; Fleming et al. 2020).

## Results and Discussion

### Bougainvillia cf. mucus morphology and phylogeny

*Bougainvillia* medusae were approximately 6 mm in height and width and had four red structures at the corners of the bell. Higher magnifications revealed that these structures were tentacle bulbs, each with 7 simple eyes (ocelli) (Fig 1). Medusae had a branching manubrium common to the genus (Fig 1). Morphological identification coupled with 18S/COI/16S alignments and phylogenetic analyses all supported the species identification as *Bougainvillia muscus* (Fig S1). A more data rich phylogenetic placement based on genes from across the genome was consistent with this identification, with *Bougainvillia* falling as sister to *Turritopsis* to form the Pseudothecata (Fig 1; Fig S2) (Mendoza-Becerril et al. 2018).

### Genome assembly

The final *Bougainvilia* genome assembly consisted of 350 contigs with an N50 of 10 Mb, max length of 26.349 Mb, and a total genome length of 375,328,287 bases (Table 1). Diverse cnidarian species including *Hydra*, *Nematostella*, *Hydractinia* and *Clytia* have 15 chromosomes (Leclère et al. 2019; Simakov et al. 2022; Zimmermann et al. 2023; Kon-Nanjo et al. 2023), so we might expect a similar number in *Bougainvillia*. While we did not recover 15 chromosomes, the number of contigs is comparable to those of the *Hydractinia* genome before scaffolding with Hi-C (Kon-Nanjo et al. 2023). The *Bougainvillia* genome was smaller than those of other hydrozoans. The *Hydractinia* genome is ∼483 Mb, the *Clytia* genome is ∼445 Mb, the *Hydra vulgaris* 105 strain is ∼819 Mb, *Hydra* AEP is ∼901Mb, and *Hydra oligactis* is ∼1274 Mb (Leclère et al. 2019; Simakov et al. 2022; Zimmermann et al. 2023; Kon-Nanjo et al. 2023; Cazet et al. 2023). The GC content of the *Bougainvillia* genome assembly was 36.21%, indicating it is AT-rich like other Hydrozoans (Chapman et al. 2010; Schnitzler et al. 2024).

**Table 1.**
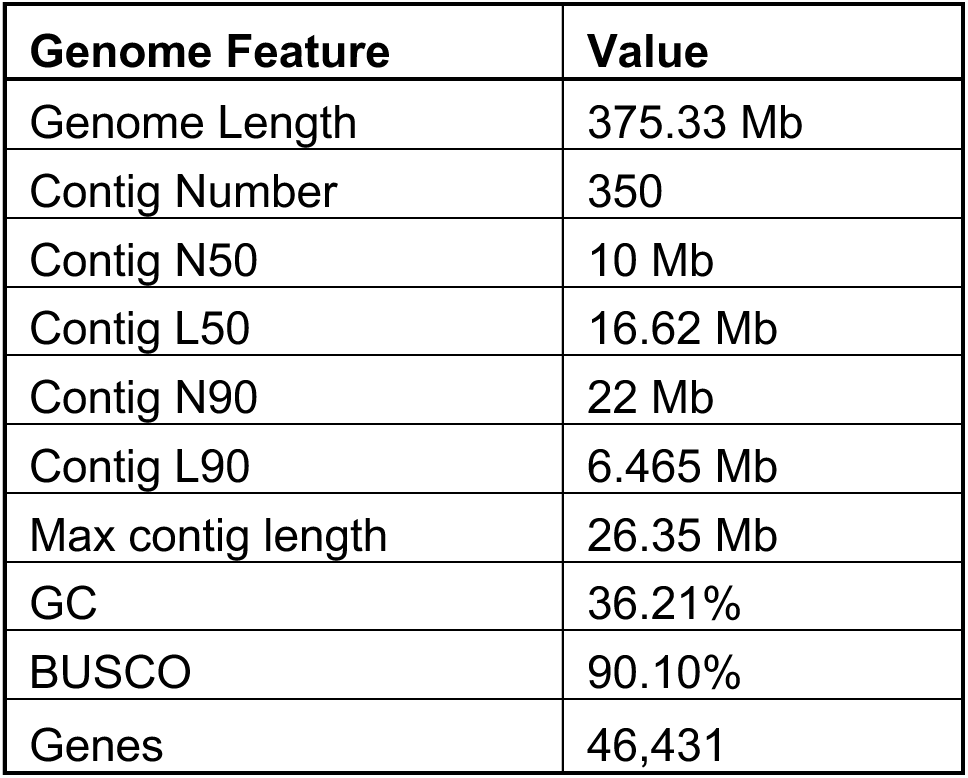
Genome Assembly Statistics.

| **Genome Feature** | **Value** |
| --- | --- |
| Genome Length | 375.33 Mb |
| Contig Number | 350 |
| Contig N50 | 10 Mb |
| Contig L50 | 16.62 Mb |
| Contig N90 | 22 Mb |
| Contig L90 | 6.465 Mb |
| Max contig length | 26.35 Mb |
| GC | 36.21% |
| BUSCO | 90.10% |
| Genes | 46,431 |

The final *Bouganvillia* assembly had a complete BUSCO score of 90.1%. Using BRAKER3 we identified 46,431 predicted protein coding genes (Table 1). This number is larger than those of the other cnidarian genomes, which have ∼20-30 thousand genes (Leclère et al. 2019; Gold et al. 2019; Zimmermann et al. 2023; Cazet et al. 2023). The number of predicted genes in our assembly is likely due to splice variants included as multiple separate gene models. To our knowledge, this is the most contiguous assembly for a hydrozoan outside of *Hydra* and *Hydractinia* (which lack eyes). Compared to eyed cnidarians, including *Aurelia* and *Turritopsis*, our genomes assembly has a smaller contig number and similar BUSCO score denoting its high quality.

### Synteny

Due to the limited number of high-quality genomes within Cnidaria, the degree of macrosynteny across the phylum remains understudied. However, previous studies in cnidaria suggest some phylogenetic signal in macrosynteny (Zimmermann et al. 2023; Kon-Nanjo et al. 2023). For example, *Nematostella* has a high degree of macrosynteny conservation with a closely related sea anemone (*Scolanthus*) but less conservation with *Hydra* and *Xenia* (CITE Zimmerman). To determine the level of macrosynteny conservation between *Bougainvillia* and two other Hydrozoans, we compared our assembly to the chromosome-level assemblies of *Hydra vulgaris* 105 strain and *Hydractinia symbiolongicarpus*. *Hydra* and *Hydractinia* have high macrosynteny conservation, with two potential translocations (Kon-Nanjo et al. 2023). Here, we found a high degree of macrosynteny conservation between *Bougainvillia* and *Hydractinia* (Fig 2A). We did not detect any obvious genome rearrangements and calculated the conservation index to be 0.863.

**Figure 2.**
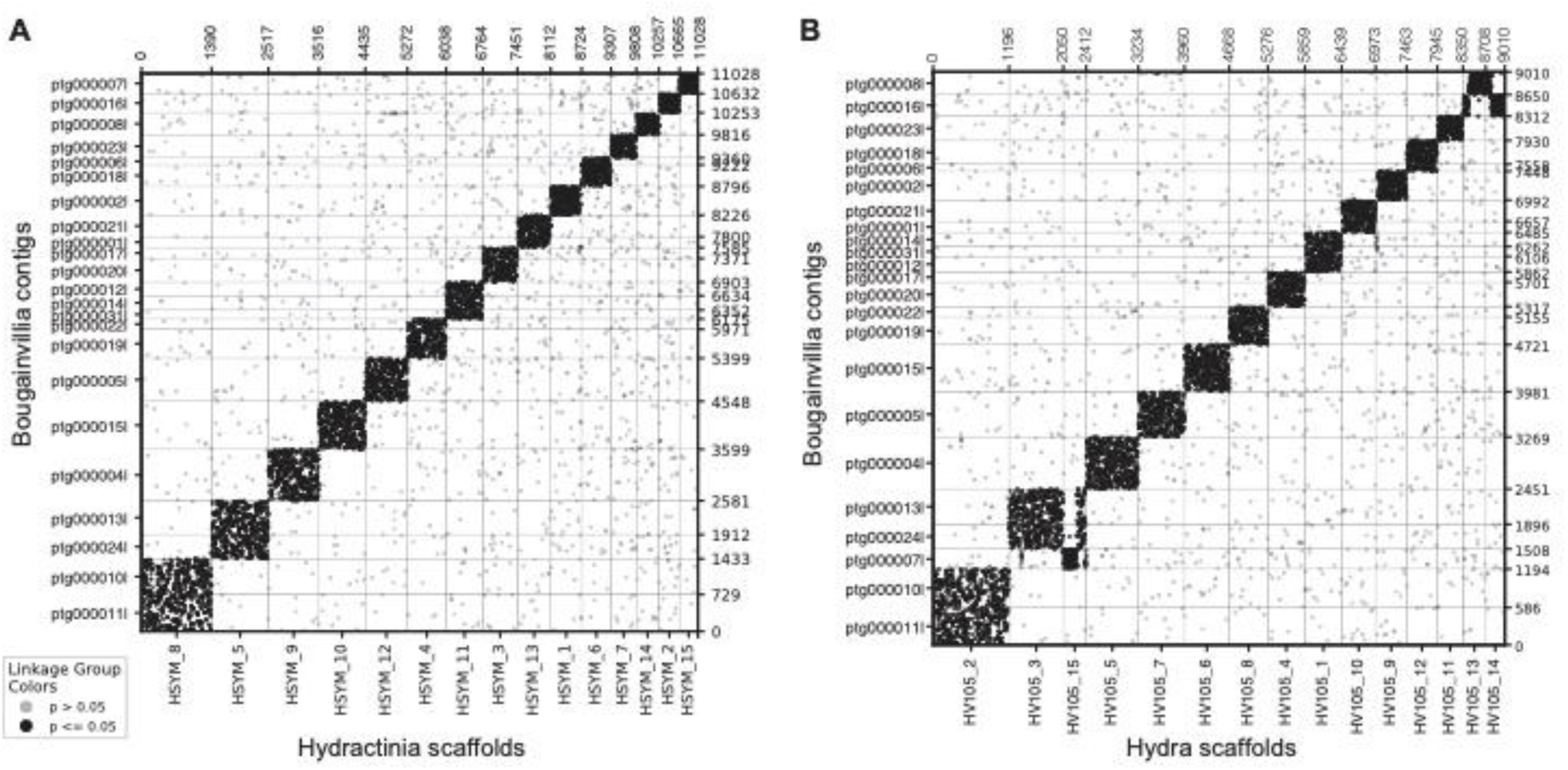
*Bougainvillia* is syntenic with other Hydrozoans. A) Oxford dot plots showing the location of 11,029 orthologous genes in 23 *Bouganvillia* contigs and 15 *Hydractinia symbiolongicarpus* scaffolds. B) Oxford dot plots showing the location of 9,011 orthologous genes in 23 *Bouganvillia* contigs and 15 *Hydra vulgaris* scaffolds.

Between *Bougainvillia* and *Hydra*, we found macrosynteny to be mostly conserved, with some potential chromosome rearrangements (Fig 2B). Firstly, there was a potential translocation between *Hydra* chromosomes 3 and 15 and *Bouganvillia* contigs ptg000013l, ptg000024l and ptg000007l (Fig 2). *Bougainvillia* contigs ptg000013l and ptg000024l were syntenic to most of *Hydra* chromosome 3 and part of chromosome 15. Meanwhile, ptg000007l was syntenic to a small part of *Hydra* chromosome 3 and syntenic to two pieces of chromosome 15. The other potential genomic rearrangement was ptg000016l which was syntenic to parts of *Hydra* chromosomes 13 and 14. These translocations are consistent with the macrosynteny analysis between *Hydra* and *Hydractinia* supporting that the translocation occurred after the divergence of *Hydra* and the *Hydractinia* and *Bougainvillia* clade. The conservation index between *Bougainvillia* and *Hydra* was 0.838. Our results suggest that *Bougainvillia* is more syntenic with *Hydractinia* than with *Hydra*, reflecting their phylogenetic relationships.

### Repetitive elements

Repetitive elements make up ∼61% of the *Hydractinia symbiolongicarpus* genome and ∼70% of the *Hydra vulgaris* genome (Kon-Nanjo et al. 2023; Cazet et al. 2023). A recent study identified active TEs driving variation in genome size between two *H. vulgaris* strains (Kon-Nanjo et al. 2024). In addition, an in-depth characterization of TEs in *Hydractinia* found the subfamily Helitron is responsible for a recent expansion of repetitive elements (Kon et al. 2024). To compare repetitive content in *Bougainvillia* to other hydrozoans, we classified the number of repetitive elements in our genome assembly. Similar to *Hydractinia*, repetitive elements made up 61.78% of the *Bougainvillia* genome, with ∼48% being unclassified or unknown (Fig 3A; Table 2). Following unclassified elements, retroelements in the LINE group made up the next largest percent of repetitive elements, 5% of the genome. Unlike in *Hydractinia*, whose sequence divergence analysis suggested two conspicuous episodes of repetitive element expansions, in *Bougainvillia* expansion of repetitive elements appears to be more continuous but also shows two peaks of repeat expansions (Fig 3B).

**Figure 3.**
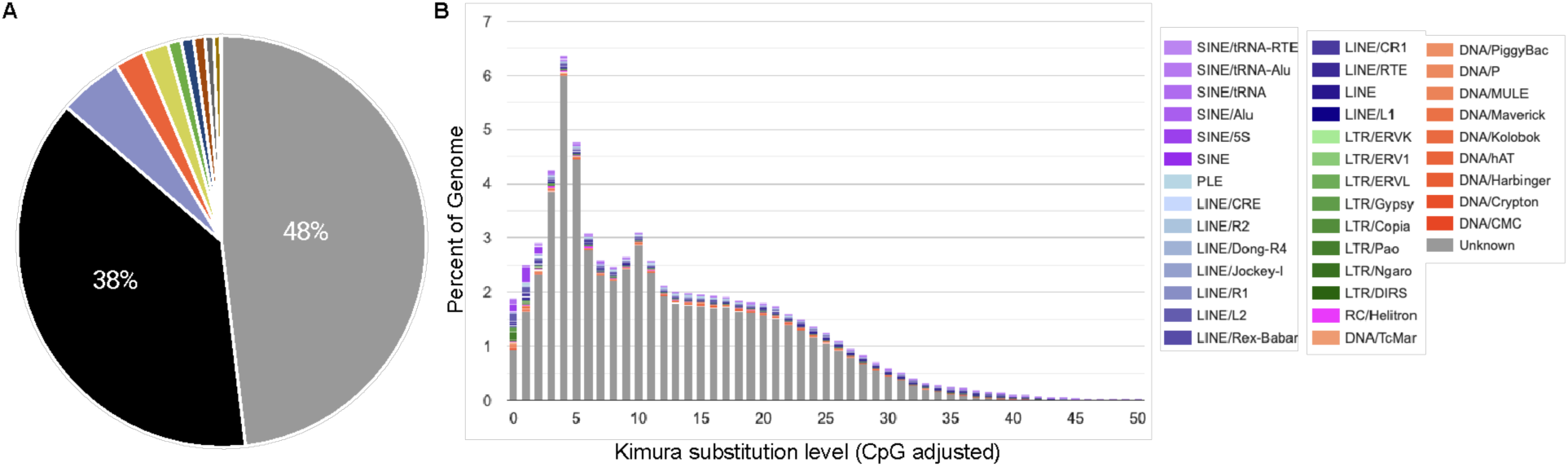
The *Bougainvillia* genome is highly repetitive. A) Pie-chart showing the percent of the genome made up of unclassified repetitive elements (gray), LINE (blue), DNA transposons (orange), small RNA (yellow), SINE (green), LTR (dark blue), Penelope (magenta), rolling circles (dark gray), simple repeats (brown), and unmasked (black). B) Repeat landscape of the *Bougainvillia* genome assembly.

**Table 2.**
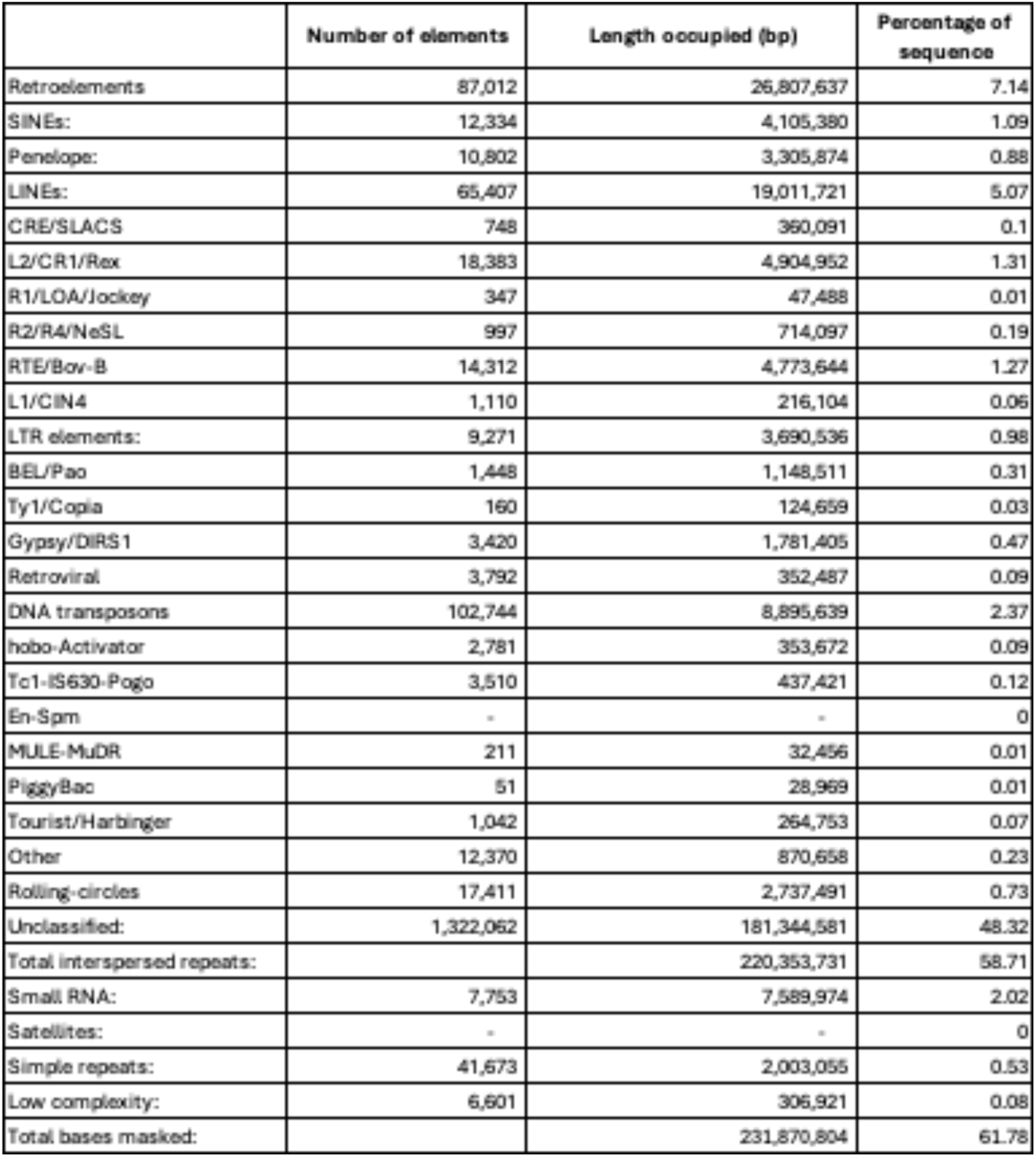
RepeatMasker output for the *Bougainvillia* genome.

### Vision-related genes

#### Opsins

Opsin genes encode proteins that are typically used to detect light. Since *Bougainvillia* has ocelli, we examined opsin gene content. We identified a total of 24 predicted opsin-like genes in *Bougainvillia* (Figure 4; Table S2). On the opsin phylogenetic tree, we found four of the genes grouped outside of the cnidops (cnidarian opsin) group and were not opsins based on additional BLAST analyses, but rather other G-protein coupled receptors (Figure 4). The remaining 20 genes were cnidops. This is comparable to the 17 opsins identified in *Tripedalia* (Liegertová et al. 2015). This is expected for medusozoa, which have a smaller diversity of opsins compared to Anthozoa (McCulloch et al. 2023).

**Figure 4.**
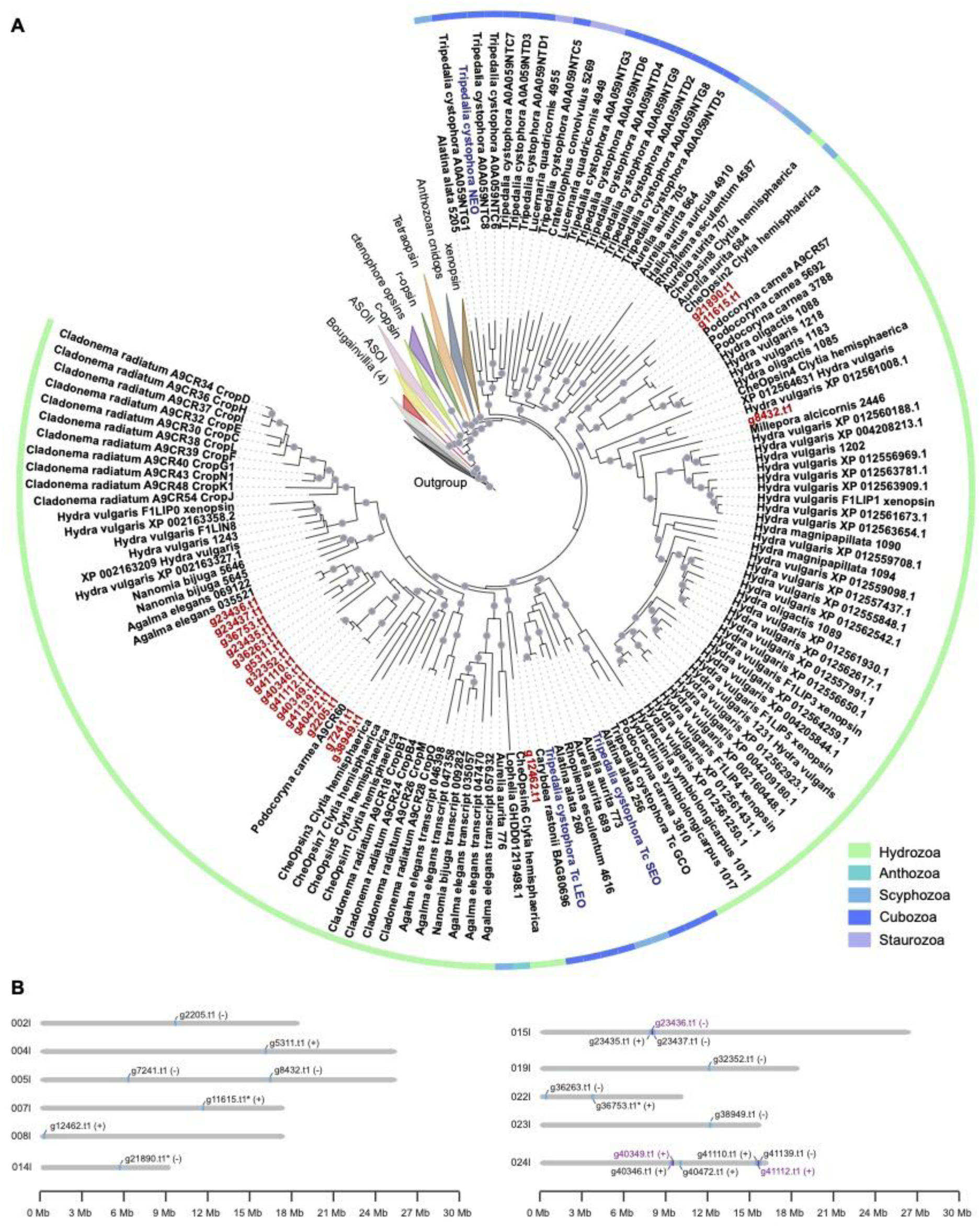
*Bouganvillia* opsin phylogeny and genome location. A) Cnidarian opsin (cnidops) phylogeny. *Bougainvillia* opsins are labeled in red. *Tripedalia* opsins known to be expressed in eyes are shown in blue. Branches that have over 80% bootstrap support are labeled with circles. B) Schematic of cnidops locations on the genome. Contigs are shown as gray bars and cnidops locations are colored and labeled (typically blue). Genes that are located close to each other are labeled in purple. Orientation is shown after the gene name in parentheses. Genes with asterisks after their name are those that have introns.

Of the 20 cnidops genes, two group together on the phylogenetic tree (*g21890.t1* and *g11615.t1*) and are closely related to *Clytia* opsin *CheOpsin2*. Another gene, *g8432.t1* groups with a *Hydra* cnidops group. One *Bougainvillia* cnidops *g12462.t1* groups together with *CheOpsin6*. The remaining cnidops form a clade together with a *Podocoryna* A9CR60 and *Clytia* CheOpsins 1,5,7,3 (Fig 4). In terms of phylogenetic position, none of the *Bougainvillia* opsin genes were orthologs of cnidops expressed in *Tripedalia* eyes (Bielecki et al. 2014; Garm et al. 2022)(Fig 4). This suggests *Bougainvillia* may be using a non-orthologous opsin for visual function, indicating probable paralog switching. The 20 *Bougainvillia* cnidops genes are distributed across 11 chromosomes and all but three, lack introns (Fig 4; Table S1). This is consistent with medusozoan opsins being intronless (Plachetzki et al. 2007; Liegertová et al. 2015). We also identified some cnidops that are located in tandem. This is the case for three *Bougainvillia* cnidops on contig ptg000015l and two pairs of genes on ptg000024l. Interestingly, the six cnidops genes on ptg000024l group together on the phylogenetic tree indicating similarity in gene sequence, including the two pairs of genes found in tandem (Fig 4). The structure and location of the *Bougainvillia* opsins suggest molecular evolution by retrotransposition and tandem duplication similar to *Hydra* and *Nematostella* opsins (Macias-Munõz et al. 2019; McCulloch et al. 2023).

#### Other vision-related genes

To investigate the molecular evolution of other candidate vision-related genes in *Bougainvillia*, we identified genes similar to those involved in eye development, phototransduction, and visual cycling from model organisms. We generated phylogenetic trees and counted the number of genes present in each gene family within the *Bougainvillia* genome, *Tripedalia cystophora* (a box jellyfish, which all have eyes) transcriptome, and *Hydra* (an eyeless hydrozoan) gene models (Table 3; Fig. S3-S23). The gene families BMP/GDF, Hedgehog, and Wnt function in visual system specification, and have undergone duplications and expansions in many animal groups, so we expected to find differences in copy number. We found several genes to be present in *Bougainvillia* and *Tripedalia* but absent in eyeless *Hydra*. These included *dpp-like*, *wnt11-like*, *wnt4-like*, and *SIX1/2-like*. Dpp -or decapentaplegic-is necessary for eye patterning in *Drosophila* and can induce ectopic expression of photoreceptor cell differentiation (Pignoni & Zipursky 1997). SIX-family genes also play a role in retinal specification in *Drosophila*. For phototransduction, G-alpha subunits, Adenlyate Cyclase (AC) and Cyclic Nucledotide Gated (CNG) channels have been identified as probable components of cnidarian phototransduction cascades (Plachetzki et al. 2007; Koyanagi et al. 2008) but diversity across clades is possible (Vöcking et al. 2022). For phototransduction and visual cycling genes, we found some potential instances of gene duplication or loss across the three cnidarians. However, except for G-alpha-s (Koyanagi et al. 2008), the specific functions of these candidate genes in vision have never been demonstrated in any cnidarian. Future studies of eye development and phototransduction in cnidarians will reveal whether these genes play functional roles in eye patterning and light-detection.

**Table 3.**
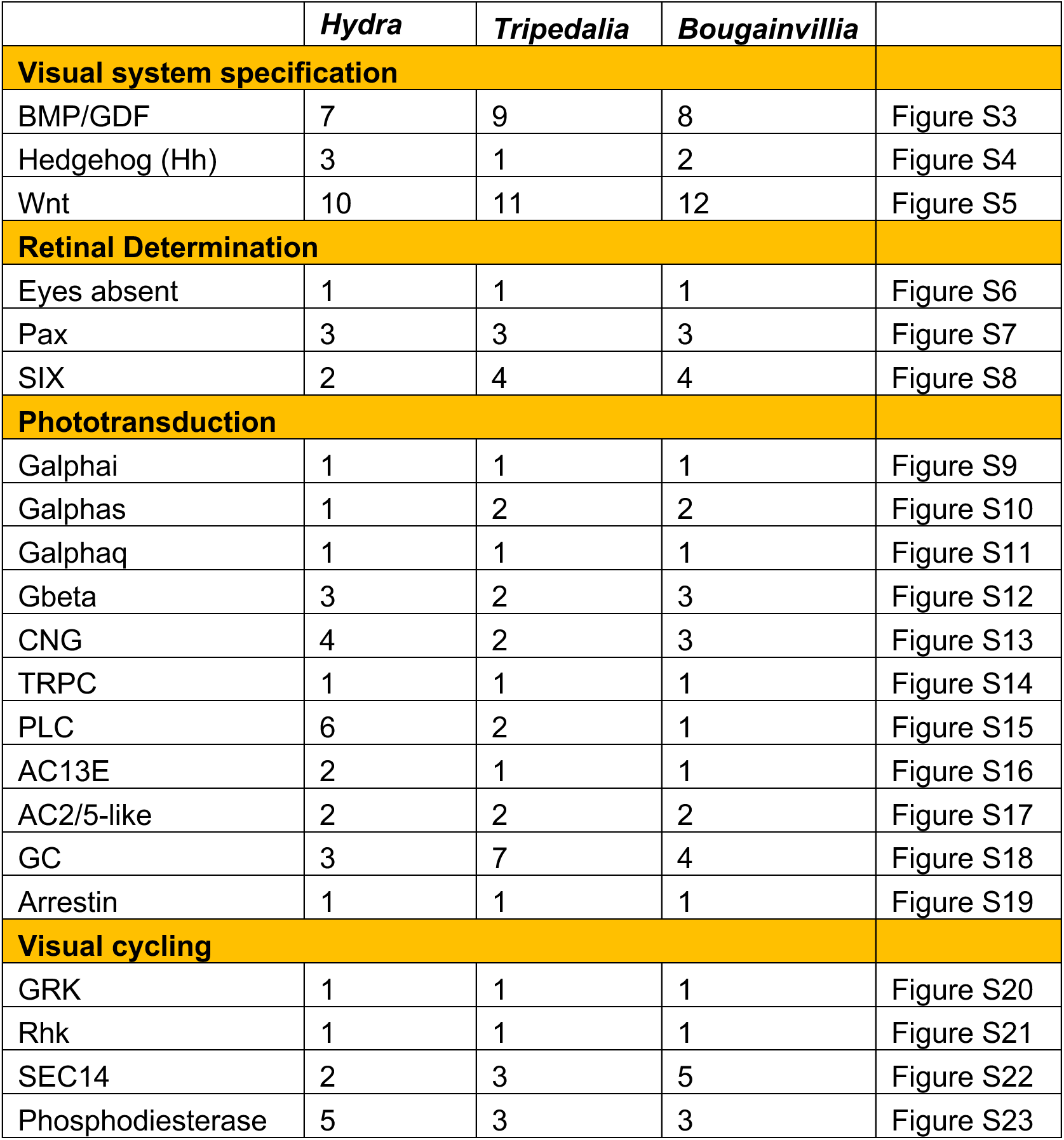
Summary of vision-related gene copy numbers.

## Conclusions

We generated the most high-quality and contiguous genome to date for any hydrozoan with a visual system. The *Bougainvillia* genome assembly consists of 350 contigs with an N50 of 10 Mb and a total genome length of 375 Mb. Though the *Bougainvillia* genome is smaller than other hydrozoans, macrosynteny is highly conserved with *Hydractinia symbiolongicarpus* and *Hydra vulgaris*, supporting synteny across hydrozoans. Similar to *Hydractinia*, ∼62% of the *Bougainvillia* genome was made up of repetitive elements. Exploration of vision-related genes identified candidate genes for future studies of eye development and light detection in cnidarians. This new hydrozoan genome is valuable for comparative studies in cnidarian and metazoan biology.

## Supporting information

Suppmental_tables_S1-2

## Data availability

Raw sequencing read data is available on SRA under accession XXXXXXXXXX. The final genome assembly and associated files are available for download at Genbank accession XXXXXX. All scripts associated with this manuscript are available on Zenodo 10.5281/zenodo.14727809. Opsin sequences were submitted to genbank accessions XXXXXX.

## Acknowledgements

The authors thank Maria Pia Miglietta and George Matsumoto for guidance in identifying the medusae to the species level and Joseph Ryan for pre-concatenated sequences and advice on a pipeline for a species-level phylogenetic tree. The authors also Anders Garm for the animal samples and everyone in the Oakley lab who assisted with animal care. This project was funded by NSF DEB-2153773 to THO and an American Philosophical Society Franklin Research Grant to AMM.

